# The *Staphylococcus aureus* iron-regulated surface determinant A (IsdA) increases SARS CoV-2 replication by modulating JAK-STAT signaling

**DOI:** 10.1101/2022.06.27.497883

**Authors:** Mariya I. Goncheva, Richard M. Gibson, Ainslie C. Shouldice, Jimmy D. Dikeakos, David E. Heinrichs

**Affiliations:** Department of Microbiology and Immunology, University of Western Ontario, London, Ontario, Canada N6A 5C1; ImPaKt Laboratory, Schulich School of Medicine and Dentistry, University of Western Ontario, London, Ontario, Canada N6A 5C1

## Abstract

The emergence and spread of Severe Acute Respiratory Syndrome Coronavirus 2 (SARS CoV-2) and the associated Coronavirus disease (COVID-19) pandemic have affected millions globally. Like other respiratory viruses, a significant complication of COVID-19 infection is secondary bacterial co-infection, which is seen in approximately 25% of severe cases. The most common organism isolated from co-infection is the Gram-positive bacterium *Staphylococcus aureus*. Here, we developed an *in vitro* co-infection model where both CoV-2 and *S. aureus* replication kinetics can be examined. We demonstrate CoV-2 infection does not alter how *S. aureus* attaches to or grows in host epithelial cells. In contrast, the presence of replicating *S. aureus* enhances the replication of CoV-2 by 10-15-fold. We identify this pro-viral activity is due to the *S. aureus* iron-regulated surface determinant A (IsdA) and this effect is mimicked across different SARS CoV-2 permissive cell lines infected with multiple viral variants. Analysis of co-infected cells demonstrated an IsdA dependent modification of host transcription. Using chemical inhibition, we determined *S. aureus* IsdA modifies host Janus Kinase – Signal Transducer and Activator of Transcription (JAK-STAT) signalling, ultimately leading to increased viral replication. These findings provide key insight into the molecular interactions that occur between host cells, CoV-2 and *S. aureus* during co-infection.

**Importance:** Bacterial co-infection is a common and significant complication of respiratory viral infection, including in patients with COVID-19, and leads to increased morbidity and mortality. The relationship between virus, bacteria and host is largely unknown, which makes it difficult to design effective treatment strategies. In the present study we created a model of co-infection between SARS CoV-2 and *Staphylococcus aureus*, the most common species identified in COVID-19 patients with co-infection. We demonstrate that the *S. aureus* protein IsdA enhances the replication of SARS CoV-2 *in vitro* by modulating host cell signal transduction pathways. The significance of this finding is in identifying a bacterial component that enhances CoV-2 pathogenesis, which could be a target for the development of co-infection specific therapy in the future. In addition, this protein can be used as a tool to decipher the mechanisms by which CoV-2 manipulates the host cell, providing a better understanding of COVID-19 virulence.

## Introduction

In December 2019, reports emerged of a pneumonia-like illness in the city of Wuhan, China. The disease was attributed to a novel coronavirus – Severe Acute Respiratory Syndrome Coronavirus 2, (SARS CoV-2, CoV-2)(1, 2) and, on March 11^th^, 2020, the WHO had declared this a global pandemic(3). CoV-2 infection, referred to as the COVID-19 disease, results in respiratory symptoms of varying severity, often including cough and fever(3). To date, there have been more than 541 million confirmed cases of COVID-19 worldwide, and over 6.3 million deaths (as of June 2022). Outside of vaccination (4, 5), treatment options for COVID-19 infections remain somewhat limited, and predominately focused on preventing death in severe, hospitalized patients.

One significant complication of respiratory viral infections, including CoV-2, is increased susceptibility to secondary bacterial infections. Although frequency of co-infection varies between reports and locations(6–8), the overall rate in the general population is roughly 5%(9). However, incidence jumps to 25-30% of hospitalized patients, and is as high as 40% in intensive care unit (ICU) patients(7, 9–11). Importantly, the mortality of CoV-2 and bacteria co-infected patients can be as high as 35%, despite the almost universal administration of antibiotics(6, 9). At present, no studies have examined the molecular interactions that occur between the two pathogens during co-infection and how that may impact disease outcome.

Data from patients infected with CoV-2, and with laboratory confirmed bacterial co-infection, show the most commonly isolated species to date has been the Gram-positive opportunistic pathogen *Staphylococcus aureus*. Prevalence of *S. aureus* varies between studies, with reports ranging from 35-70% of isolates being *S. aureus*, with both methicillin sensitive (MSSA) and methicillin resistant (MRSA) isolates reported in most studies(6, 9–11). However, despite the abundance of *S. aureus* co-infection in COVID-19 patients, little is known about how or if the virus and bacteria affect each other, and what effect this may have on pathogenesis.

In previous studies with other respiratory viruses, *S. aureus* has also been shown as a frequent cause of secondary bacterial co-infection(12, 13). In the case of influenza A virus (IAV), several studies have demonstrated co-infection results in more severe immune system dysregulation(13–15), including depletion of alveolar macrophages. Non-immune mediated interactions have also been characterised at the molecular level(16–18) - IAV infection results in increased adhesion of both *S. aureus* and *Streptococcus pneumoniae* to epithelial cells and, in the case of *S. aureus*, increased intracellular bacterial replication(18). Conversely, the *S. aureus* protein lipase 1 enhances IAV replication in primary cells through the positive modulation of infectious particle release(19). Based on the similarity of clinical presentation and co-infection frequency, we reasoned events during co-infection will be similar between IAV – *S. aureus* co-infection and CoV-2 *– S. aureus* co-infection. In the present study, we investigated the interplay between CoV-2 and *S. aureus* and demonstrate that *S. aureus* enhances the replication of CoV-2 *in vitro* through the bacterial iron regulated surface determinant protein A (IsdA). The expression of IsdA leads to a modification of Janus Kinase – Signal Transducer and Activator of Transcription (JAK-STAT) signalling in CoV-2 infected cells, which positively regulates viral replication.

## Results

### *S. aureus* enhances the replication of CoV-2 in epithelial cells

Previous reports have indicated that infection with IAV results in increased adhesion to and replication of *S. aureus* in epithelial cells(18). Based on these findings, we sought to determine if infection of cells with CoV-2 also affects adhesion of *S. aureus*. To do this, we developed an *in vitro* co-infection model where the replication kinetics of both pathogens could be quantified (outlined in Figure 1A). We used the Wuhan isolate of CoV-2, the MRSA strain USA300 LAC and a bacterial mutant lacking fibronectin binding proteins (FnbAB), the latter proteins being required for invasion of *S. aureus* into epithelial cells(20, 21). We observed that *S. aureus* efficiently adheres to Vero E6 cells (Figure 1B), whereas a preceding CoV-2 infection did not impact bacterial adhesion (Figure 1B, Supplementary Figure 1A, 1B). Further, *S. aureus* was able to invade into Vero E6 cells, and this was dependent on the presence of FnbAB (Figure 1B), as previously shown for other epithelial cells(20–22). However, no differences in invasion rates were observed between cells alone and CoV-2 infected cells (Figure 1B, Supplementary Figure 1C). Since bacterial invasion was equivalent between uninfected and CoV-2 infected cells, we next tested if preceding viral infection impacts the subsequent rate of intracellular bacterial replication. To test this, we examined bacterial numbers at 6h, 8h and 20h post invasion. As shown in Figure 1C, bacterial replication occurs, as demonstrated by increased bacterial numbers over time. However, the rate of bacterial replication, and the absolute level to which the bacteria grew, did not differ between cells alone and CoV-2 infected cells. When CoV-2 titre was determined, we observed that the presence of *S. aureus* increased the amount of infectious virus particles, at both 12h and 24h post infection (Figure 1D). Taken together, these data indicate that CoV-2 infection does not overtly impact the interaction of *S. aureus* with epithelial cells, but that *S. aureus* enhances CoV-2 replication in Vero E6 epithelial cells.

**Figure 1.**
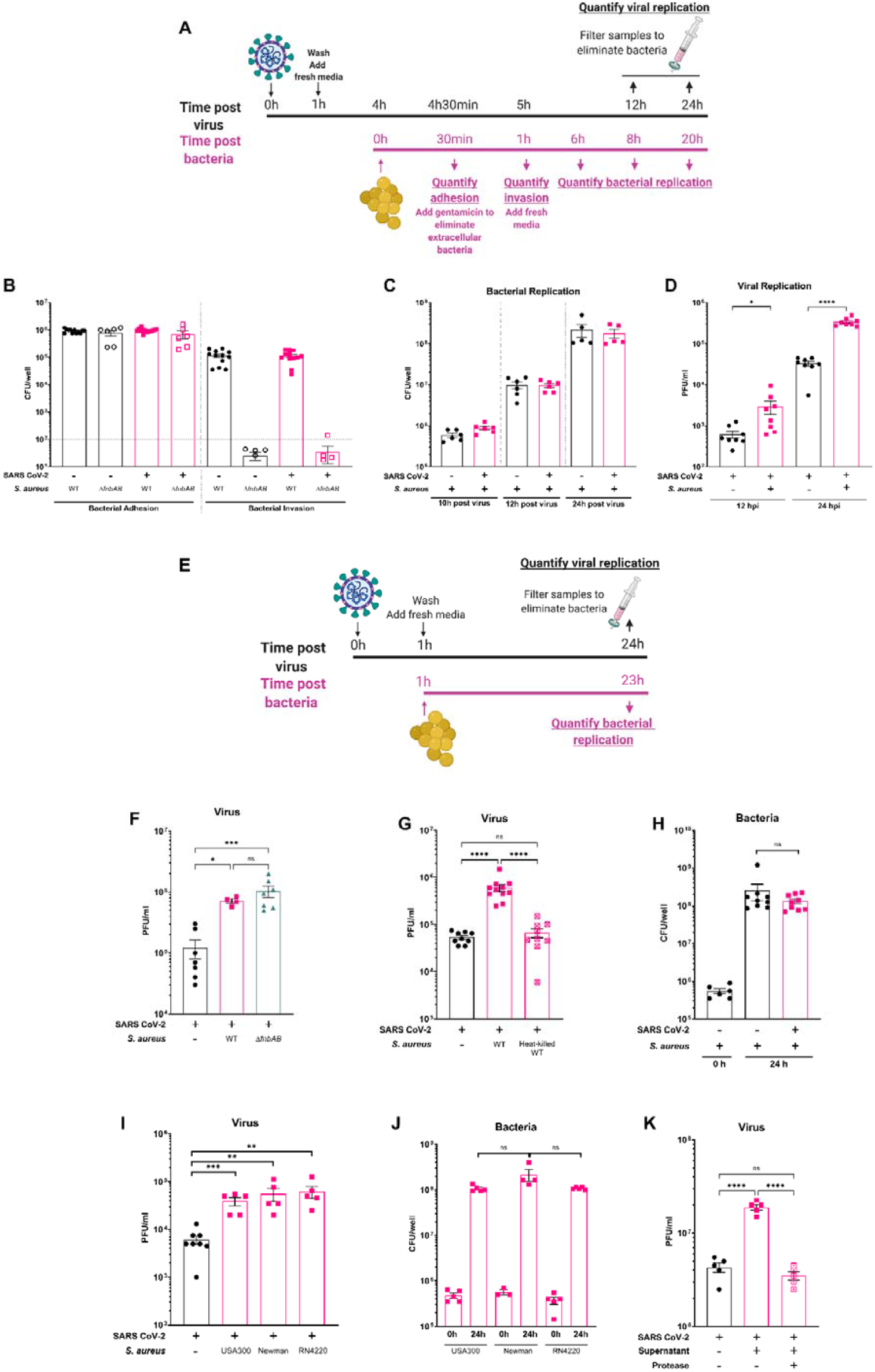
*S. aureus* enhances CoV-2 replication in Vero E6 cells. **A** - schematic representation of experimental procedures in the *in vitro* co-infection model. **B** - Vero E6 cells were mock infected or infected with CoV-2 at MOI of 1 for 1h, and at 4h, *S. aureus* was added to the cells at an MOI of 10. Bacterial adhesion was assessed after 30 min, by lysing Vero E6 cells and enumerating the total number of CFU. Representative wells were treated with 150 µg/mL of gentamicin for 30 min, cells were washed 2 times with PBS and then lysed for bacterial enumeration. Horizontal dotted line shows limit of accurate detection. **C** - Cells were treated as in B, except post washing, cells were overlayed with 1mL of SF DMEM. At indicated times, DMEM was removed, and cells lysed for enumeration of bacteria. **D** - Cells were treated as in C, except at indicated times, DMEM was harvested, samples were centrifuged for 1 min at 13000 x *g*, and the resulting supernatant was passed through a 0.22 µm filter before being used to determine viral titre in a standard plaque assay on Vero E6 cells. **E** – schematic representation of the modified experimental model used for the remainder of the study. **F** - Vero E6 cells were infected with CoV-2 at MOI of 1 for 1h and overlayed with 1mL of SF DMEM. 1×10^5^ CFU of WT or *ΔfnbAB S. aureus* were added to the culture media. 24h later, DMEM was harvested, samples were centrifuged for 1 min at 13000 x *g*, and the resulting supernatant was passed through a 0.22 µm filter before determination of viral titre. **G** - cells were infected as in F, except 1×10^5^ CFU of either *S. aureus* or heat killed *S. aureus* were added. At 24h, samples were harvested and viral titre determined as in F. **H** - the pelleted samples from G were re-suspended in 500 µL PBS + 0.1 % Triton X-100 and bacterial CFU was enumerated. **I** – Cells were infected as in **F**, and 1×10^5^ CFU of *S. aureus* strains USA300, Newman or RN4220 added. **J** – Pelleted sampled from **I** were processed as in **H. K** - Cells were infected with CoV-2 at MOI of 1 and 50 µL of either TSB, bacterial supernatant, or protease treated bacterial supernatant were added to the SF DMEM. Infections were harvested 24h later and viral titre determined. For all experiments, viral titre was determined through plaque assay on Vero E6 cells. Data shown are mean ±SEM of 3-5 independent experiments. In some experiments (B, C, D, G, H), each replicate included 2 independent bacterial cultures. Statistical analysis - unpaired student’s t test. *p<0.05, ** p<0.01, *** p<0.01, **** p<0.001.

Given that the model employed in Figure 1A demonstrated that *S. aureus* enhances virus replication, it’s tempting to hypothesize that CoV-2 and *S. aureus* have replicated in the exact same cell. To determine if this is the case, we simplified the model and excluded bacterial invasion (Figure 1E). We added 1×10^5^ CFUs of WT or the invasion incapable *ΔfnbAB S. aureus* mutant, the approximate amount detected to become intracellular in Vero E6 cells (see Figure 1B), to the culture medium of infected cells. Using this method, we observed the same, ∼10-fold increase in viral titre (Figure 1F), demonstrating that bacterial invasion is not needed, and the addition of *S. aureus* to the extracellular environment is sufficient for increased virus replication to occur.

Next, we sought to determine if the presence of *S. aureus* cells is enough to enhance CoV-2 growth, and if the bacteria even need to be alive. We examined virus replication in the presence of WT *S. aureus*, or the equivalent number of heat-killed bacteria. While the WT bacteria enhanced virus growth by ∼10-fold, no effect was observed in the presence of heat-killed bacteria (Figure 1G). Concurrently, we observed that the WT bacteria grew by over 2-log in the 24h period of the experiment (Figure 1H). Furthermore, when we examined additional strains of *S. aureus*, including an avirulent laboratory strain RN4220, we saw equivalent levels of pro-viral activity (Figure 1I) and bacterial replication (Figure 1J), demonstrating this effect is not restricted to USA300. Based on these findings, we reasoned the pro-viral activity is due to either an increase in the number of bacteria over the 24h, or to a factor produced during bacterial growth. To test the latter option, we added cell free supernatant of stationary phase WT *S. aureus* to CoV-2 infected cells. This supernatant contains a large number or proteins, including enzymes, virulence factors, and other by-products of bacterial replication. Indeed, we observed that the supernatant alone was sufficient to increase CoV-2 replication (Figure 1K), albeit not to levels observed with the whole bacteria. To determine if the bacterial pro-viral factor is a protein, we treated the supernatant with the broad-spectrum protease trypsin, which resulted in degradation of observable polypeptides (Supplementary Figure 2). This treatment also eliminated the pro-viral phenotype (Figure 1K), demonstrating that CoV-2 replication is enhanced by a *S. aureus* protein or proteins that are present in the bacterial supernatant.

### The *S. aureus* iron regulated surface determinant A (IsdA) mediates the bacterial pro-viral activity

Considering we had observed the pro-viral phenotype with both virulent and avirulent *S. aureus* strains and shown cell-free supernatant was sufficient for this activity, we hypothesised a protein or proteins that are secreted or released by the *S aureus* cell are responsible. To test this, we employed bacterial mutants, lacking one or more of the key *S. aureus* proteins normally found in the bacterial supernatant and added them to virus infected cells (as in Figure 1E). As shown in Figure 2A, all these mutants retained pro-viral activity, with the exception of the mutant in sortase A (*srtA*). Sortase A is an endopeptidase that covalently links the group of proteins known as “cell-wall anchored” (CWA) proteins to the peptidoglycan of a bacterial cell, resulting in their display on the cell surface(23, 24). However, many of these proteins are subsequently digested by proteases and released in the bacterial supernatant. To confirm the role of sortase A, we complemented the bacterial mutant by providing the full-length genes *in trans* on a plasmid. As shown in Figure 2B, complementation successfully restored the pro-viral phenotype, without impacting bacterial replication of these strains (Figure 2C).

**Figure 2.**
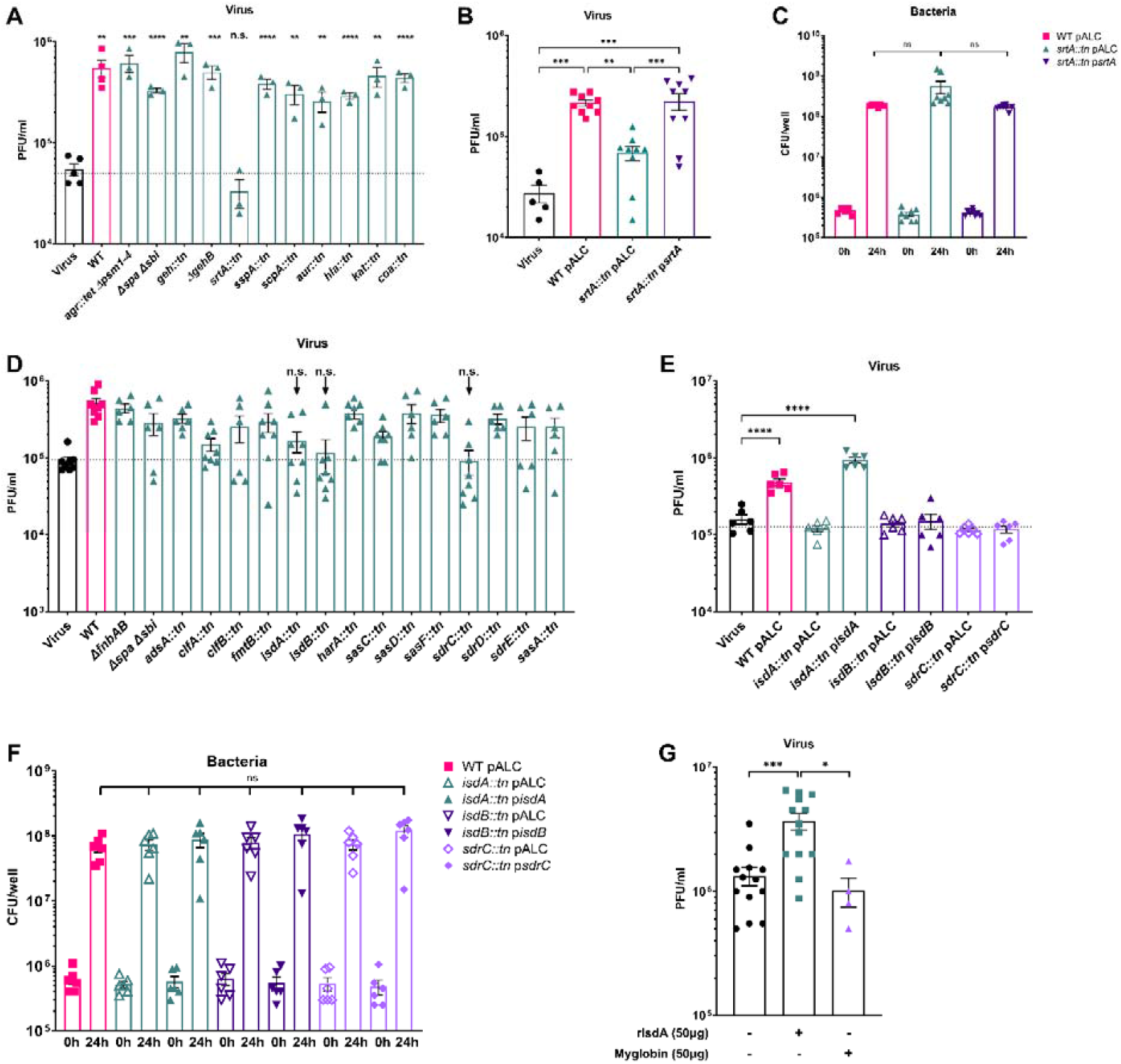
The *S. aureus* iron regulated surface determinant A (IsdA) mediates pro- viral activity. **A** - Vero E6 cells were infected with CoV-2 at MOI of 1 for 1h and overlayed with 1ml of SF DMEM. 1×10^5^ CFU of WT or indicated mutants of *S. aureus* were added to the culture media. Infections were harvested at 24h and viral titre was determined. Data shown are mean SEM of 3 independent experiments. **B** – Cells were infected as in **A**, but with strains carrying empty vector controls or complementing plasmids. Data shown are mean SEM of 3 independent experiments, with 3 replicates per experiment. **C** - the pelleted samples from B were re-suspended in 500 µL PBS + 0.1 % Triton X-100 and bacterial CFU was enumerated. **D –** cells were infected as in A, but different *S. aureus* mutants were added. Data shown are mean SEM of 3 independent experiments, with 2 replicates per experiment. **E –** cells were infected as in A, but with strains carrying empty vector controls or complementing plasmids. Data shown are mean SEM of 3 independent experiments, with 2 replicates per experiment. **F –** samples from E were processed as in C. **G** – Vero E6 cells were infected as in A, but 50µg of recombinant IsdA, myglobin or equal of buffer (50mM HEPES, pH 7.2) were added. Samples were harvested at 24h and viral titre determined. For all experiments, viral titre was determined through plaque assay on Vero E6 cells. Statistical analysis – B, E, - one-way ANOVA with multiple comparisons between all samples, and Turkey’s post-test. A, C, D, F and G - unpaired student’s t test. *p<0.05, ** p<0.01, *** p<0.01, **** p<0.001.

In the absence of sortase A, CWA proteins are still produced, but are instead directly secreted into the bacterial supernatant(23, 24); we also noticed that only partial elimination of pro-viral activity is seen with the *srtA* mutant. Taken together, these findings suggest that virus replication is enhanced by one of the proteins anchored by sortase A, rather than the sortase itself. To test this hypothesis, we took individual mutants of each of the 18 proteins encoded in the strain USA300 (except *fnbAB* and *spa sbi*, where double mutants were used), and tested them for pro-viral activity. As shown in Figure 2D, all but three of these mutants retained pro-viral activity – *isdA, isdB* and *sdrC*. To determine the role of these 3 genes, we complemented each mutant by providing the full-length genes *in trans* on a plasmid and re-tested them for pro-viral activity. Only provision of *isdA* restored the phenotype (Figure 2E), even though all strains grew to the same level (Figure 2F). Of note, we also observed that complementation of *isdA* resulted in increased secretion of the protein in the bacterial supernatant and restoration of IsdA detection on the bacterial cell surface (Supplementary Figure 3). However, we were unable to confirm the same for IsdB and SdrC due to the unavailability of antibodies. Therefore, the role of these two proteins should not be ruled out as potential further factors enhancing CoV-2.

As we had previously observed secreted bacterial proteins were sufficient for pro-viral activity, we sought to determine whether recombinant IsdA can enhance CoV-2 replication on its own. Indeed, we observed that when rIsdA was added to virus infected cells a modest, but still significant increase in viral titre was observed (Figure 2G). This effect was not present when we employed myoglobin, an eukaryotic protein that, like IsdA, also carries a heme molecule. Nevertheless, the effect of the recombinant IsdA was not as pronounced as the whole bacteria, suggesting IsdA shed from the bacteria is different, presumably at least in the presence of the peptidoglycan it is anchored to. As such, we continued characterisation of IsdA’s effect on CoV-2 replication using whole bacteria.

Having identified at least one bacterial factor responsible for the pro-viral activity on Vero E6 cells, we next sought to determine the scope of the phenotype. We tested bacterial strains on the human lung epithelial A549 cells, which were stably engineered to express ACE2 and TRMPSS2. Although the presence of WT *S. aureus* did not enhance CoV-2 replication, we observed higher levels of host cell damage with this cell line, likely due to the activity of many host-restricted *S. aureus* toxins(25) (Supplementary Figure 3A). Indeed, when we tested a modified strain that produces only minimal levels of toxins (*agt::tet* Δ*psm1-4* has reduced toxin production due to inactivation of a major virulence regulator(26)) we did see higher viral titre, compared to cells treated with WT *S. aureus* (Supplementary Figure 4A). Nevertheless, overexpression of *isdA* in the complemented strain of either background resulted in ∼10 fold more virus being produced (Supplementary Figure 4A), despite equal bacterial replication levels (Supplementary Figure 4B). Although infection of primary human bronchial-epithelial cells resulted in low levels of infectious virus production by 24h, we nevertheless still demonstrate an increase in titre when the *isdA* overexpressing strain of *S. aureus* was present (Supplementary Figure 4C, 4D).

Furthermore, similar to observations made with the WT (Wuhan) CoV-2 virus, the Delta variant was also enhanced by *S. aureus* in an IsdA dependent manner during infection of Vero E6 or A549 ACE2 TRMPSS2 cells (Supplementary Figure 4E, 4F). Indeed, we also detected pro-viral activity for the recombinant IsdA protein for the Delta variant, in both Vero E6 and A549 ACE2 TRMPSS2 cells (Supplementary Figure 5). Overall, these data demonstrate that *S. aureus* pro-viral activity is mediated through IsdA, it is conserved over different cell types and viral variants, and is retained in a recombinant form of IsdA.

### *S. aureus* IsdA induces specific host transcriptional changes

In order to determine how *S. aureus* and IsdA affect CoV-2, we performed an RNAseq analysis of Vero E6 cells treated with the WT, *isdA::tn* or *isdA::tn* p*isdA* bacteria (Figure 3A). To ease data interpretation and eliminate any virus specific effects on the cells, these experiments were performed in the absence of virus (Figure 3A). Analysis of differentially regulated genes (DEGs) of samples with bacteria indicated that despite hundreds of DEGs detected, only 11 genes were in common between the comparisons of WT vs *isdA::tn* and WT vs *isdA::tn* p*isdA* (Supplementary File 2, Figure 3B, Table 1). This suggests that the presence of *isdA* has a specific transcriptional effect on only these genes, and other changes can be attributed to different *S. aureus* proteins.

**Figure 3.**
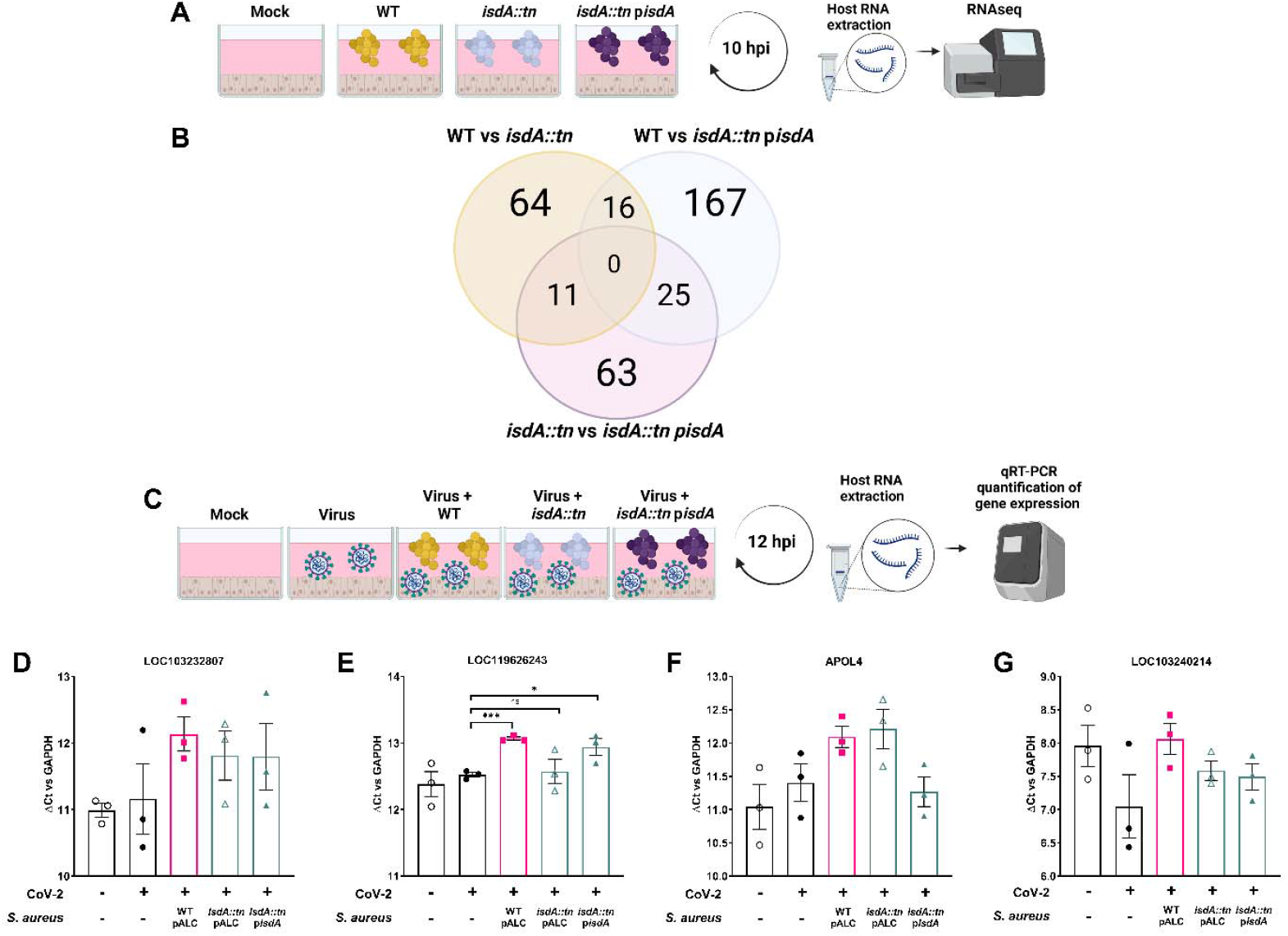
*S. aureus* expressing IsdA specifically modulates host cell transcript levels. **A –** Schematic representation of the experimental model used for RNAseq sample generation. **B** – Numbers of differentially expressed genes between cells treated with different bacterial mutants. **C** – Schematic representation of the experimental model used for RT-PCR sample generation. **D - G** – Vero E6 cells were infected as shown in C, total RNA extracted and RT-PCR was performed for the indicated genes. All Ct values were normalised to GAPDH levels of the sample. Data shown are mean ± SEM of 3 independent experiments. Statistical analysis –unpaired student’s t test. *p<0.05, ** p<0.01, *** p<0.01.

**Table 1.** Host genes that were differentially expressed when exposed to *S. aureus* expressing IsdA (i.e. present in the comparison of both WT vs *isdA::tn* and *isdA::tn* vs *isdA::tn pisdA*). A log2 fold change cut-off of >1 and < -1 was used.

| Locus | Description | Homologous Human Gene | WT vs *isdA::tn* |  | *isdA::tn* vs *isdA::tn pisdA* |  |
| --- | --- | --- | --- | --- | --- | --- |
|  |  |  | Log2 Fold change | p value | Log2 Fold change | p value |
| LOC103232807 | rho GTPase-activating protein 4 | ARHGAP4 | -5.85 | 0.026 | -5.61 | 0.033 |
| LOC103235349 | zinc finger and SCAN domain-containing protein 5B-like | ZSCAN5B | -2.97 | 0.0004 | -1.96 | 0.014 |
| LOC119622462 | uncharacterized |  | -2.17 | 0.001 | -2.02 | 0.001 |
| LOC119626243 | collagen alpha-1(I) chain-like | JAK1 | -1.69 | 0.0002 | -1.10 | 0.008 |
| LOC103229191 | uncharacterized | GPHN | -1.58 | 0.007 | -1.20 | 0.033 |
| APOL4 | apolipoprotein L4 | APOL4 | -1.34 | 0.015 | -1.48 | 0.007 |
| LOC103240214 | probable E3 ubiquitin-protein ligase HECTD2 | HECTD2 | -1.25 | 0.026 | -2.21 | 0.0003 |
| LOC119624411 | uncharacterized | AOPEP | -1.06 | 0.048 | -1.18 | 0.027 |
| LOC119622958 | uncharacterized | N/A | -1.06 | 0.04 | -1.11 | 0.029 |
| LOC103241337 | PSME3-interacting protein-like | PSME3 | -1.04 | 0.048 | -1.07 | 0.043 |
| PIWIL1 | piwi like RNA-mediated gene silencing 1 | PIWIL1 | 1.78 | 0.008 | 1.64 | 0.011 |

As our RNAseq analysis did not include CoV-2 infected cells, we sought to determine if any of the DEG we identified as modified by the presence of *isdA* also display transcriptional changes during co-infection. Accordingly, we next infected cells with CoV-2 or CoV-2 and *S. aureus* (as detailed in Figure 3C) and extracted host RNA. We quantified transcript levels by RT-PCR for the genes with highest level of change seen by RNAseq and excluded uncharacterised proteins. We also tested LOC103235349 and PIWIL1, however no transcript levels were detected. As shown in Figure 3D-G, we saw the presence of *S. aureus* changed the expression of all 4 of the genes tested. However, these changes were *isdA* dependent in only one case, where we saw WT and p*isdA* carrying bacteria decrease the expression of LOC119626243, while the *isdA::tn* mutant showed expression levels similar to the virus alone (Figure 3E). The locus is annotated as collagen alpha-1(I) chain-like protein, and a more detailed bioinformatic analysis identified sequence similarity to the human Janus Kinase 1 (JAK1) gene. Altogether these data suggest that the presence of IsdA on *S. aureus* cells triggers a specific response in host cells during both bacterial infection and CoV-2 co-infection, and a JAK1 like gene is one of the key transcripts affected.

### *S. aureus* IsdA modulates JAK2-STAT3 signalling to enhance CoV-2 replication

Given that we observed significant changes in the transcription of a gene with homology to JAK1 during co-infection, we decided to investigate the expression of the four JAK genes in Vero E6 cells. We examined cells at 12h post virus infection, both in the presence and absence of bacteria, and assessed transcription through RT-PCR. As shown in Figure 4 (A-D), we observed that co-infection resulted in a small increase in JAK1 transcripts in the presence of WT *S. aureus*, but no transcriptional changes were observed for JAK2, JAK3 or TYK2. Furthermore, the effect on JAK1 was not specific to the presence of *isdA*.

However, given that JAK-STAT signalling occurs through protein expression and/or post translational modifications such as phosphorylation, it is unlikely that significant differences would be seen at the transcript level. Therefore, we further examined the role of the JAK-STAT pathway at the protein/function level. Effectively, we chose to use chemical inhibition, as the antibody availability for the cell line used is limited or cross-species reactivity of the antibodies has not been tested. Pan-JAK inhibitors (CP 690550 citrate, Pyridone 6) at 5µM decreased CoV-2 replication both in the presence and absence of *S. aureus*, making them unsuitable for co-infection investigations (data not shown). However, the inhibitor SD1008, which targets the signal transduction between JAK2 and STAT3(26), eliminated the pro-viral effect of *S. aureus* without impacting viral replication alone (Figure 4E). Furthermore, SD1008 inhibition was specific to *S. aureus* expressing *isdA*, suggesting this is at least one of the mechanism/s through which IsdA enhances CoV-2 replication. Importantly, the presence of the 1µM SD1008 did not impact the ability of *S. aureus* to replicate, demonstrating the decrease in viral titre was not due to absence of bacterial growth (Figure 4F). In addition, we observed an equivalent inhibition of the *S. aureus* pro-viral effect when SD1008 was added to cells infected with the Delta variant of CoV-2 (Figure 4 G, H). These data indicate the activity of the inhibitor, just as we had seen with whole bacteria and recombinant IsdA, is not restricted to a specific viral variant. Taken together, these findings suggest inhibition of JAK2 and/or STAT3 activation is at least partially responsible for the IsdA mediated pro-viral effect of *S. aureus*.

**Figure 4.**
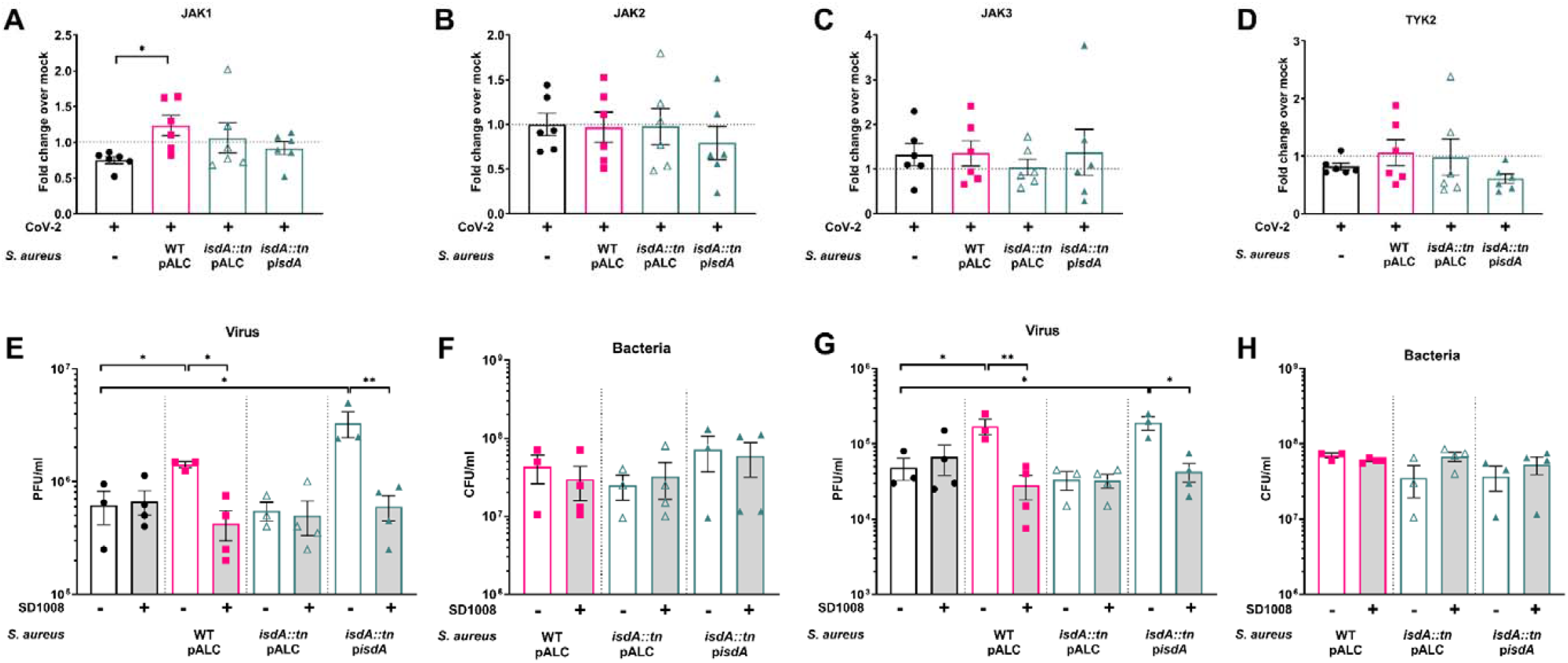
*S. aureus* IsdA affects JAK-STAT signalling to promote virus replication. **A-D** Vero E6 cells were infected with CoV-2 at an I of 1 for 1h and overlayed with 1mL of SF DMEM. 1×10^5^ CFU of WT or indicated mutants of *S. aureus* were added to the culture media. At hpi, total RNA was extracted, and RT-PCR was performed for the indicated genes. All Ct values were normalised to GAPDH levels of the mple. Data shown are mean ± SEM of 5 independent experiments. **E-H** Cells were infected as in **A**, and 1 µM of SD1008 or DMSO were ded concurrently with the bacteria. At 24hpi, viral titre was determined by plaque assay. The pelleted bacteria were re-suspended in 500 µL S + 0.1 % Triton X-100 and bacterial CFU was enumerated. Data shown are mean ± SEM of 3-4 independent experiments. Statistical alysis –unpaired student’s t test. *p<0.05, ** p<0.01.

## Discussion

The emergence and spread of SARS CoV-2, and the subsequent COVID-19 pandemic have demonstrated the devastating effect respiratory viral pathogens can have on human life and health. Morbidity and mortality of respiratory viruses, both during seasonal outbreaks and pandemics, are complicated by increased susceptibility of patients to secondary bacterial co-infection. *S. aureus* has historically been one of the most common organisms identified in co-infection, especially in the case of influenza; similarly, worldwide data indicates *S. aureus* is also the most prevalent bacterial species in CoV-2 co-infection patients(9). Strikingly, co-infection with *S. aureus* can increase mortality from ∼0.8% with CoV-2 alone, to as high as 35% during co-infection(9). In COVID-19 patients, work has demonstrated that acute CoV-2 infection and the associated increase in cytokine levels decreases the bacterial killing capacity of neutrophils and monocytes, therefore contributing to the development of bacterial co-infection(27). Immune system dysregulation has also been shown to play a significant role in the pathogenesis of co-infection with influenza, where the immune response skewing from the initial viral replication creates a pro-bacterial environment in the host(13–15). The importance of these extreme responses is undeniable for the inability of the host to clear the infection. However, molecular interactions between viruses and bacteria during co-infection are relatively unstudied in comparison, due to the requirement for complex models and/or specific cell types. Here, we demonstrate the identification of the first bacterial protein that can manipulate the host cell to favour CoV-2 replication. The broad effect of IsdA on different CoV-2 variants, coupled with the universal presence of this gene in *S. aureus* isolates suggests the impact of IsdA on co-infection pathogenesis can be significant.

Using a co-infection system where the replication kinetics of both pathogens can be measured, we demonstrated *S. aureus* enhances the ability of CoV-2 to replicate (Figure 1). These data are similar to observations made with IAV, where *S. aureus* was also pro-viral(18, 28). In contrast, the presence of the virus provided no benefit to bacterial attachment or replication. This differs from IAV, where viral infection increased the ability of *S. aureus* and *S. pneumoniae* to adhere to and invade into host cells(18). The variation in results between IAV and CoV-2 suggests virus-specific interactions occur with co-infecting bacteria, even when clinical frequency and presentation are similar.

Utilizing bacterial mutants, we were able to identify CoV-2 replication was impacted by the bacterial protein IsdA (Figure 2). In *S. aureus*, IsdA is part of a network of Isd proteins anchored into the bacterial cell wall, which serves to transport heme iron into the bacterial cell(29, 30). However, some IsdA and/or IsdA attached to peptidoglycan is released from the cells as the peptidoglycan layer is renewed. Interestingly, IsdA has been shown to allow *S. aureus* adherence to squamous epithelial cells in the nares(31, 32), and the widespread expression of the protein during infection has made it an attractive target for inclusion as a vaccine component(33). To our knowledge, this is the first report of IsdA manipulating host cell transcription and signal transduction. The effect this has on viral replication is likely an inadvertent consequence of targeting pathways that could also be beneficial to the bacteria. IsdA is now the second *S. aureus* protein shown to impact viral replication through host modulation. However, the previous report of *S. aureus* lipase 1 and IAV identified an effect specific to primary fibroblast cells(19). In contrast, we see IsdA impacts cellular transcription and/or signal transduction pathways and can be seen in both primary and immortalised cells. The observed differences could be due to different approaches, as the effect of lipase 1 on cell transcription remains unknown. Indeed, we also have not examined how IsdA effects CoV-2 replication in fibroblast cells, or whether a cumulative effect would be observed if both proteins are present, and a suitable cell type is used. Nevertheless, the possibility that IsdA effects are specific to CoV-2 cannot be ruled out, and further studies are necessary to test IsdA’s potential effect on other viruses.

Our data indicate *S. aureus* IsdA manipulates the JAK/STAT signalling in the cell, which ultimately results in the observed pro-viral effect (Figure 3, Figure 4). JAK/STAT signalling serves to transmit a signal from the cell surface, usually when a molecule is bound, resulting in phosphorylation of JAK, subsequent phosphorylation of a STAT protein, and eventually transcriptional changes in the cell. As JAK-STAT signalling is also triggered by binding of cytokines and chemokines, it is a major path for induction of intrinsic cellular immunity, including the activation of Interferon stimulated genes(34). Given that immune dysregulation and “cytokine storms” are a significant contribution to COVID-19 mortality, the concept of bacterial co-infection further skewing this response can go a long way to explain the high mortality rates observed in co-infected patients.

The central role of JAK/STAT signalling in responding to and inducing immune signals has made it a target for many virus induced manipulations. Indeed, CoV-2 was recently demonstrated to downregulate JAK signalling and inhibit the phosphorylation of JAK1 and Tyk2(35). Assessment of patients has also shown increased STAT1 and phospho-STAT1 levels in peripheral monocytes(36). It would be of great interest to compare if these responses are further elevated if bacterial co-infection is present. The original SARS CoV virus also decreases STAT1 phosphorylation through the action of the non-structural protein 1(37), and dephosphorylation of STAT3 is seen during infection of Vero E6 cells(38).

Work on other RNA viruses has also shown IAV decreases the phosphorylation of STAT1(39) and avian infectious bronchitis virus Nsp14 facilitated degradation of JAK1 and impaired the nuclear translocation of STAT1(40). Furthermore, a study of IAV – *S. aureus* co-infection also demonstrated *S. aureus* prevents the dimerization of STAT1 and STAT2, which results in increased IAV replication(28). It will be interesting to determine if this STAT1-STAT2 dimerization block is mediated by IsdA, or if *S. aureus* produces more than one protein that can manipulate this pathway. Indeed, the question also remains of the level of change in JAK and STAT expression and phosphorylation by IsdA, and what benefit this provides to the bacteria. Nevertheless, we believe IsdA can serve as a tool to further understand how CoV-2 manipulates the cell during replication, and whether these mechanisms can be targeted for therapeutic purposes.

Overall, our work demonstrates a new link between CoV-2 and co-infecting bacteria, and indeed one of the first reports of a direct molecular interactions between a coronavirus and bacteria. The identification of IsdA as a pro-viral factor shows that there is still much that is unknown about how specific bacterial proteins interact with respiratory viruses. Further characterisation of such events can provide a simultaneous two-fold advancement of knowledge, in both co-infection events, and as new tools to study the biology of viruses.

## Materials and methods

### Tissue culture

African Green Monkey kidney Vero E6 cells were purchased from the ATCC and maintained in Dulbecco’s modified Eagle’s medium (DMEM) with 10% (v/v) fetal bovine serum (FBS) at 37°C, 5% CO_2_ and passaged twice a week. A549 ACE2 TRMPSS2 cells were a kind gift from Dr Matthew Miller (McMaster University, Canada), and were maintained in DMEM + 10% w/v FBS, 700µg/ml G418, 800µg/ml hygromycin B and 1%w/v L-Glutamine) at 37°C, 5% CO_2_ and passaged twice a week. Primary human bronchio/tracheal epithelial cells were purchased from the ATCC and maintained in airway epithelial medium, as recommended by ATCC. Primary cells were not used past passage 7.

### Bacterial growth

Bacterial strains and plasmids used in this study are listed in Supplementary Table 1. *E. coli* was grown in Luria-Bertani (LB) broth and *S. aureus* was grown in tryptic soy broth (TSB) at 37°C, shaken at 200 rpm, unless otherwise stated. Where appropriate, media were supplemented with erythromycin (3 µg/mL), chloramphenicol (12 µg/mL), lincomycin (10 µg/mL), kanamycin (50 µg/mL), tetracycline (3µg/mL) or ampicillin (100 µg/mL). Solid media were supplemented with 1.5% (w/v) Bacto agar.

### PCR and construct generation

*S. aureus* strain USA300 LAC, cured of the 27-kb plasmid that confers antibiotic resistance, was used as the WT strain for mutant generation, unless otherwise stated. Primers used in this study are listed in Supplementary Table 2. Transposon insertion mutants were obtained from the Nebraska transposon mutant library. For complementation, the full-length genes were amplified, ligated into pALC2073 and transformed into *E. coli*. All plasmids were passaged through RN4220, prior to transfer to the strain of interest.

### Western blot

For detection of secreted bacterial proteins, bacteria were grown overnight in TSB, bacteria were pelleted and the supernatant (equal to OD_600_ of 8) was used for a trichloroacetic acid (TCA) precipitation. Briefly, equal volumes of bacterial supernatant and 20% (w/v) TCA were mixed and incubated at 4°C for 3h. Samples were pelleted at 21 000 *x* g for 15 min, washed twice with ice-cold 70% ethanol and allowed to dry overnight. Pellets were re-suspended in 40 µL Laemmli buffer, boiled at 95°C for 10 min and 15 µL loaded on a 12% SDS-PAGE gel. Samples were run at 150V for 90 min, and transferred on a nitrocellulose membrane using a TransBlotter Turbo (Biorad) standard settings. Membranes were blocked in 5% (w/v) skimmed milk in PBS + 0.1% Tween-20 (PBST) overnight at 4°C. Primary antibody was added at 1 in 500 dilution in blocking buffer for 2h at RT, followed by 3 washes with PBST. Secondary antibody (donkey anti-rabbit IRDye 800) was added at 1 in 20 000 dilution in PBST for 1h, followed by 3 washes with PBST. Membranes were imaged on a LiCor scanner.

### Immunofluorescence

Bacteria were grown overnight in TSB, pelleted (equal to OD_600_ of 4) and fixed in 4% paraformaldehyde (PFA) for 20 min. Cells were then washed twice with PBS and incubated with tetramethylrhodamine (TMR) labelled wheatgermagglutinin (WGA) (2 µg/mL) in PBS for 1h. Cells were washed twice with PBS and blocked in 5% (w/v) bovine serum albumin for 2h at RT. Primary antibody was added at 1 in 500 dilution in blocking buffer for 2h at RT, followed by 3 washes with PBS. Secondary antibody (goat anti Rabbit AlexaFluor 488) was added at 1 µg/mL in PBS for 1h, followed by 3 washed with PBS. Cells were then re-suspended in 50 µL PBS and allowed to dry on coverslips. Coverslips were then mounted using Prolong Diamond and imaged on a Zeiss LSM880 confocal microscope.

### Recombinant protein purification

Full length isdA was generated by amplification of the gene (without the N terminal signal sequence and the C terminal region following the LPXTG anchoring motif) and ligation into pET28A+. The plasmid was then transformed into *E. coli* BL21 (DE3) and recombinant protein production was induced. Briefly, 0.5 L cultures were grown in LB with 100 µg/mL kanamycin until OD_600_ of 0.6-0.8. Cultures were then induced with 1mM IPTG for 16h at RT, pelleted and frozen at -20°C. When required, pellets were defrosted, re-suspended in 50 mL lysis buffer (50 mM NaH_2_PO_4_, 300 mM NaCl, 10 mM imidazole, pH 8.0) with complete protease inhibitor (Roche, UK), passed through a One-Shot cell disruptor (Constant systems, Northants, UK) at 30 kPsi, centrifuged at 4000x *g* for 30 min and passed through a 0.45µm filter. Proteins were purified by immobilised metal affinity chromatography (IMAC) with a FF Crude Ni-NTA column, using a gradient of 0 – 100 % elution buffer (50 mM NaH_2_PO_4_, 300 mM NaCl, 300 mM imidazole, pH 8) and dialysed in 50mM HEPES, pH 7.2. Relative protein concentration was determined using a Bradford assay.

### Virus growth and quantification

SARS CoV-2 isolates were acquired from BEI. SARS-Related Coronavirus 2 -- Isolate USA-WA1/2020 (Wuhan isolate) was deposited by the Centers for Disease Control and Prevention and obtained through BEI Resources, NIAID, NIH: SARS-Related Coronavirus 2, Isolate USA-WA1/2020, NR-52281. The Delta VOC strain used was SARS-CoV-2, B.1.617.2 variant NR-55611. All experiments with live virus were performed under Biosafety Level 3 conditions in Western University’s ImPaKt facility. For generation of virus stocks, confluent Vero E6 cells were washed with PBS, infected with passage 1 virus in 5mL of serum-free DMEM (SFM) for 1h at 37°C, 5% CO_2,_ with gentle rocking every 10 min. The inoculum was then removed, cells washed with PBS, and overlayed with DMEM + 2% FBS and incubated for 72h at 37°C, 5% CO_2._ The culture supernatant was collected, centrifuged at 500 x *g* for 10 min and aliquots were stored at -80°C. For virus quantification, standard plaque assays were performed on Vero E6 cells. Briefly, confluent monolayers of cells in 6 well tissue culture plates were washed with PBS and infected with 400 µL of virus dilutions in SFM. Plates were incubated for 1h at 37°C, 5% CO_2,_ with gentle rocking every 10 min. The inoculum was removed, and cells were overlayed with 2ml/well of a 1:1 mixture of 2.4% Avicel and 2 x Plaque overlay (91% 2xMEM, 4% FBS, 1% Pen/strep, 1% 1M HEPES, 0.5% GlutaMAX). Plates were incubated for 72h at 37°C, 5% CO_2,_ after which cells were fixed by the addition of 2mL of 10% Neutral Buffered Formalin for 30 min at RT. The fixative was washed, and cells stained with Crystal violet (80% water, 20% Methanol, 1% (w/v) crystal violet) for 15 min at RT. The stain was washed away with water, plates allowed to dry, and plaques counted.

### Adhesion, Invasion and bacterial replication in epithelial cells

For all experiments involving bacterial adhesion and invasion, confluent Vero E6 cells in 12 well tissue culture plates were used. Cells were maintained in DMEM + 10% (v/v) FBS until the day of infection. On the day of infection, cells were washed twice with PBS and infected with CoV-2 at MOI of 1 in 100 µL per well for 1h at 37°C, 5% CO_2,_ with gentle rocking every 10 min. The inoculum was then removed, cells washed with PBS and overlayed with 1mL of SFM. At indicated times post virus infection, bacteria were added at an MOI of 10.

Bacterial strains of interest were grown O/N in TSB, with appropriate antibiotics. Bacteria were then sub-cultured at OD_600_ of 0.1 and grown in TSB, with appropriate antibiotics, to OD_600_ of 0.6. Cells were then pelleted, washed twice with DMEM and re-suspended in DMEM to a density of 2×10^7^ CFU/mL. 50 µL of that suspension were added to a well of Vero E6 cells that were either uninfected or infected with CoV-2 containing 700 µL of SFM. Plates were pelleted at 1000 rpm for 1 min and incubated at 37°C, 5% CO_2_ for 30 min. For quantification of adhesion, media was then removed, cells lysed with 500 µl of PBS + 0.1% (v/v) Triton - X100 and plated for CFU. The remaining wells were then treated with 150 µg/mL gentamicin for 30 min at 37°C, 5% CO_2_, extensively washed to remove the gentamicin, and kept in SFM for the desired duration of the infection, as indicated in the text. At specific times post infection, media was removed, and cells lysed in PBS + 0.1% (v/v) Triton - X100, scraped from the well, and plated for CFU.

### Virus infection with extracellular bacteria or recombinant protein

For infections where bacteria were added extracellularly, virus infection was performed as above (see Adhesion section) at an MOI of 1. Bacterial strains of interest were grown O/N in TSB, with appropriate antibiotics. Cells were then pelleted, washed twice with DMEM and re-suspended in DMEM to a density of 2×10^7^ CFU/mL. 5 µL of that suspension was added to the 1mL of SF DMEM present in virus infected wells. At 24h, the media was harvested, pelleted at 13 000 x *g* for 1 min the resulting supernatant was passed through a 13mm diameter 0.22µm filter. Cells were lysed with 500 µL of PBS + 0.1% (v/v) Triton - X100, and the lysate was added to the pellet of extracellular bacteria, before plating for total CFU. For infections with heat killed bacteria, cells were treated as above, the 2×10^7^ CFU/mL solution was incubated at 85°C for 30 min, and 5µL of that suspension was added to virus infected cells. For infections with bacterial supernatant, WT *S. aureus* were grown O/N in TSB, OD_600_ was normalised to 4 and 50µL were added to virus infected cells. For protease treatment, supernatant was processed as above, 25 µg of TPCK Trypsin were added for 1h min at 37°C, followed by heat inactivation at 95°C for 20 min. 50µL were added to virus infected cells. For infections with recombinant protein, virus infection was done as above, and indicated concentrations of recombinant protein were added to the 1mL of SF DMEM present in virus infected wells. For infections in the presence of inhibitors, 1µM of SD1008, 5 µM CP 690550 citrate or Pyridone 6 or DMSO were added to cells immediately post inoculum removal, followed by the addition of *S. aureus* as above.

### RNA extraction and RNA sequencing

For RNAseq experiments, 12 well plates of Vero E6 cells were infected, or mock infected, with CoV-2 at an MOI of 1 and then treated with 1×10^5^ CFU *S. aureus* or respective mutants, as indicated in the respective figures. Cells were then washed 3 times with PBS and lifted with 500µL/well of Cell Protect Reagent (Qiagen) for 5 min. Samples from multiple wells were pooled (11 for RNAseq, 4-6 for RNA isolation and RT-PCR), pelleted at 1200 x *g* for 5 min and stored at -80°C overnight. The cell pellet was lysed with 500 µL of PBS + 0.1% (v/v) Triton - X100 and RNA extracted using the QIAGEN RNAEasy kit, as per the manufacture’s instructions. DNAse treatment was performed with TURBO DNA Free kit (Invitrogen) for 2 × 30 min, followed by inactivation with the supplied buffer. RNA sequencing was performed by the Microbial Genome Sequencing Center in Pittsburgh, PA. Data analysis, including read mapping and differential expression were performed by the Microbial Genome Sequencing Center in Pittsburgh, PA. Briefly quality control and adapter trimming was performed with bcl2fastq. Read mapping was performed with RSEM(41). Read counts loaded into R(42) and were normalized using edgeR’s (43) Trimmed Mean of M values (TMM) algorithm. Subsequent values were then converted to counts per million (cpm). Differential expression analysis was performed using edgeR’s Quasi-Linear F-Test (qlfTest) functionality against treatment groups.

### qPCR

1 µg of RNA was used in a reverse transcriptase reaction, as per the manufacturer’s instructions (Agilent Technologies). qRT-PCR reactions were set up in 15µL volumes, using 0.75µL of cDNA, using SYBRgreen master mix (BioRad) and run on a Roche Rotor-Gene 6000 machine. Expression was normalized to GAPDH. Primers used are shown in Table 3.

## Acknowledgements

Work in the DEH and JD laboratories was supported by Canadian Institutes for Health Research Operating grants. This work was partially supported by a Western University Catalyst grant (49635) to DEH, MG and JD, and a British society for antimicrobial chemotherapy grant to DEH and MG (BSAC-COVID-99). Much of the work in this study was performed in a CL3 laboratory, which was funded by grants from the Canada Foundation for Innovation, The Ontario Research Fund, and the Schulich School of Medicine and Dentistry, University of Western Ontario.

